# Auxin is metabolized through kynurenine in *Hypericum perforatum* L

**DOI:** 10.64898/2026.05.18.726114

**Authors:** Daniel Gaudet, Alisha Greene, Susan J. Murch, Lauren A.E. Erland

## Abstract

Recent studies have demonstrated the presence of kynurenine (KYN) and kynurenic acid (KYNA) in several plant species, but the metabolic function of these metabolites remains undefined. We hypothesized that KYN and KYNA are metabolites of auxin and play a role in plant morphogenesis. To test our hypothesis, we developed a plant tissue-culture-based bioassay using *Hypericum perforatum* (St. John’s wort; SJW), a model system for auxin and indoleamine metabolism and pharmacological inhibitors (PF-04859989, RO-61-8048, and KMO inhibitor II, JM6) of human kynurenine pathways enzymes. SJW is an interesting model system because explants root in the absence of plant growth regulators but supplementation of the culture media with 10 μM IAA induces a callus response without *de novo* root organogenesis. Supplementation of the culture media with 10 μM KYN increased root number and internodal length relative to basal media. We used a previously validated high-resolution mass spectrometry analytical method to quantify KYN, KYNA, and 3-hydroxyanthranilic acid (3-HAA). KYN, KYNA and 3-HAA were quantified in roots and shoots of SJW grown on basal media. Supplementation of the culture media with 10 μM KYN increased the concentration of KYN, KYNA and 3-HAA in roots and shoots. Treatment with 10 μM IAA increased KYN and 3-HAA concentration in shoots. Three pharmaceutical candidates that are kynurenine pathway inhibitors in humans were taken up into the tissues from the culture media and increased KYN content as compared to basal control. Together, these data propose a role for KYN in IAA metabolism, shoot and root organogenesis.

**Highlights:**

- Kynurenine metabolites are detected and accumulate in *H. perforatum* tissue culture
- IAA redirects metabolism towards accumulation of KYN and 3-HAA in shoots
- Exogenous KYN promotes KYNA accumulation
- Pharmacological inhibition alters kynurenine pathway metabolite profiles in a tissue-specific manner
- Kynurenine and IAA differentially regulate root development

## 1. Introduction

Kynurenine (KYN) kynurenic acid (KYNA) and 3-hydroxyanthranilic acid (3-HAA), are metabolites of tryptophan that have been well characterized in animals as a major route of tryptophan catabolism (Fig.1) (Badawy, 2017). In early studies to understand niacin biosynthesis in orchid embryos, radiolabel from tryptophan was recovered in KYN, indicating a similar metabolic mechanism (Cooper et al., 1982). Kynurenine pathway metabolites have been detected across multiple species (Gaudet et al., 2026; Russo et al., 2022; Turska et al., 2022; Turski et al., 2011, 2009) (Table S1). It has been proposed that KYN interacts with auxin signaling in Arabidopsis (He et al., 2011). KYN and auxin are metabolites of tryptophan (Fig 1). Tryptophan is metabolized by TRYPTOPHAN AMINOTRANSFERASE OF ARABIDOPSIS (TAA1) and related TAR enzymes (Mashiguchi et al., 2011). Kynurenine acts as a competitive inhibitor of TAA1/TAR activity potentially affecting auxin metabolism in plants (He et al., 2011). In plant-associated microbial systems, auxin was metabolized to anthranilic acid by symbiotic microbes (Conway et al., 2022). Our recent data indicated that KYN and related metabolites vary with growing conditions in St. John’s wort (*Hypericum perforatum* L.) (Gaudet et al., 2026).

**Figure 1.**
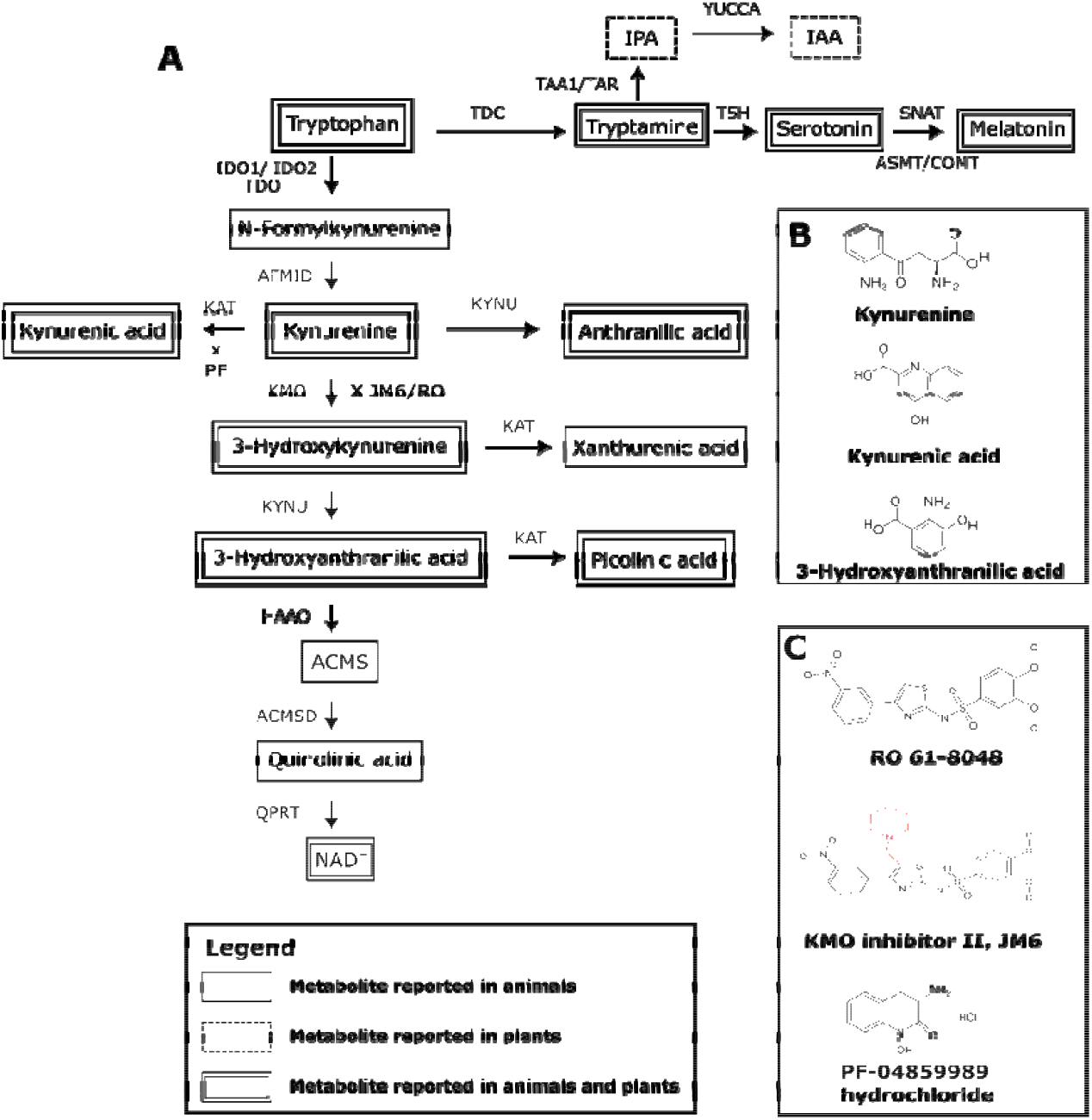
Overview of the kynurenine pathway and associated metabolites and inhibitors. (A) Schematic representation of major tryptophan (Trp)-derived metabolic pathways, including the kynurenine pathway (center), the indole-3-pyruvic acid (IPA)–indole-3-acetic acid (IAA) pathway, and the tryptamine– serotonin–melatonin branch (top). Solid, dashed, and double boxes indicate metabolites reported in animals, plants, or both, respectively. Enzymes are indicated at each step: IDO1/IDO2 (indoleamine 2,3-dioxygenase), TDO (tryptophan 2,3-dioxygenase), AFMID (arylformamidase), KAT (kynurenine aminotransferase), TDC (tryptophan decarboxylase), TAA1/TAR (tryptophan aminotransferase), KYNU (kynureninase), KMO (kynurenine 3-monooxygenase), HAAO (3-hydroxyanthranilate 3,4-dioxygenase), ACMSD (α-amino-β-carboxymuconate-ε-semialdehyde decarboxylase), and QPRT (quinolinate phosphoribosyltransferase). Inhibitor targets are indicated at the corresponding steps: JM6 and RO 61-8048 inhibit KMO, and PF-04859989 inhibits KAT. (B) Chemical structures of the kynurenine pathway metabolites quantified in this study: kynurenine, kynurenic acid (KYNA), and 3-hydroxyanthranilic acid (3-HAA). (C) Chemical structures of the inhibitors used in this study. Core structural differences between JM6 and RO 61-8048 are highlighted in red.

St. John’s wort (SJW) is a medicinal plant (Culpeper, 1816), an invasive species (Crompton et al., 1988) and a model organism for the study of indoleamine metabolism in plants (Murch et al., 2001, 2000a, 2000b; Murch and Saxena, 2004). Cell differentiation, regeneration, organogenesis and totipotency in SJW respond to concentration and localization of the indoleamine plant growth regulators serotonin and melatonin (Erland et al., 2019, 2018, 2015). Pharmaceuticals such as methylphenidate (Ritalin) and fluoxetine (Prozac) have been applied to understand biochemical mechanisms and transport of serotonin and melatonin in SJW (Murch et al., 2001; Murch and Saxena, 2004). Pharmacological inhibitors of the KYN pathway have been developed as potential drug targets in humans (Fig. 1; Jacobs et al., 2017). RO-61-8048 is a high-affinity inhibitor of kynurenine 3-monooxygenase (KMO) in mammalian systems, while JM6 is a structural derivative containing an additional 1-methylpiperidine moiety to the thiazole ring to facilitate drug manufacture (Jacobs et al., 2017). The compound (R)-3-amino-1-hydroxy-3,4-dihydroquinolin-2(1H)-one (PF-04859989) was identified by a high-throughput screen of the Pfizer compound library as a high-affinity inhibitor of human kynurenine aminotransferase (KAT; Jacobs et al., 2017). To the best of our knowledge, these compounds have not previously been evaluated for roles in plant physiology.

Our overall objective was to develop a bioassay system for investigations of KYN metabolism in plants. We hypothesized that KYN controls morphogenesis in SJW. To test this hypothesis, we evaluated the effects of exogenous indole-3-acetic acid (IAA) and KYN on shoot and root organogenesis in vitro. We quantified KYN, KYNA and 3-HAA using our previously validated analytical method (Gaudet et al. 2026). We also investigated the potential of inhibitors of the two primary mammalian enzymes for KYN catabolism (PF-04859989, RO-61-8048 and KMO Inhibitor II, JM6) as mediators of plant physiology. Our data show significant metabolic cross-talk between auxin and KYN in the expressions of totipotency in plants.

## 2. Materials and Methods

### 2.1 Plant materials and growth conditions

*Hypericum perforatum* L. stock cultures were maintained as perpetual axenic cultures as previously described (Murch and Saxena, 2006). Cultures were grown on Murashige and Skoog (MS) medium (Murashige and Skoog, 1962) supplemented with Gamborg B5 vitamins (Phytotech Labs (Gamborg et al., 1968), USA), 3% sucrose, and 2.25 g L^−1^ Phytagel (Sigma-Aldrich), pH 5.7. All cultures were grown in a controlled environment growth room at 25 ± 0.5 °C under a 12 h photoperiod provided by full-spectrum LED lighting (EcoGrowTech, Kelowna, Canada). Light conditions at culture height were ≈ 6450 lux (≈ 60 µmol m^−2^ s^−1^), with a correlated color temperature of 3642 K.

### 2.2 Experimental media and treatment conditions

Explants were excised from 6-week-old *H. perforatum* shoot cultures. To ensure uniformity, tissues were excised three nodes below the shoot apical meristem and consisted of a stem segment containing approximately 7–8 mm of internode tissue on either side of a single node with an attached pair of leaves. Explants were aseptically transferred to experimental media at time zero (T = 0) and maintained under the same controlled environment growth conditions. Experimental media were prepared using MS basal salts supplemented with B5 vitamins, 3% (w/v) sucrose, and 2.25 g L^−1^ Phytagel. Medium pH was adjusted to 5.7 prior to autoclaving at 121 °C and 15 psi for 20 min resulting in a basal control medium hereafter referred to as MSO. Following autoclaving, MSO medium was cooled to approximately 55 °C before addition of sterile-filtered IAA (Sigma-Aldrich), KYN (Millipore Sigma), inhibitors KMO Inhibitor II (JM6), PF-04859989 (PF), and RO-61-8048 (RO) (Millipore Sigma) alone or in combination. Sterile stock solutions of IAA were prepared by accurate weighing into a volumetric flask, dissolving in 1.0 mL of 0.5 N NaOH and bringing to final volume with ultrapure water. Kynurenine was weighed, dissolved in 1.0 mL methanol and diluted to final concentration with ultrapure water. All KYN pathway inhibitors were prepared in dimethyl sulfoxide (DMSO). Stock solutions were sterilized by passage through a 30-mm, 0.22-µm nylon syringe filter (Avantor/VWR) and added aseptically to cooled MS-based medium. Kynurenines and inhibitors were tested either as stand-alone treatments (0 µM or 10 µM IAA, KYN, JM6, PF-04859989, RO-61-8048) or as co-applications in which inhibitors were combined with IAA. The addition of these stock solutions resulted in final solvent concentrations below 0.1% (v/v) in all treatments. Solvent matched controls containing DMSO at equivalent concentrations were included as part of the physiological assay validation. Following incorporation of the sterile stock solutions, 1 mL of medium was pipetted into sterile 10 × 75 mm borosilicate glass culture tubes (Fisher Scientific). After the medium had cooled and solidified, explants were placed into the tubes and sealed with Parafilm.

### 2.3 Physiological measurements

Physiological responses were observed and samples were collected after 35 days. Root initiation was scored as a binary trait, with rooting defined as the presence of at least one visible root emerging from the explant. Maximum root length was measured for all rooted plantlets by gently removing the explant from the culture tube and placing it flat against a ruler; the longest root on each plantlet was recorded. Shoot height was measured for all plantlets, regardless of rooting status, as the distance from the base of the explant to the shoot apex using a ruler. Representative images were taken at 35 days using a dissecting microscope (Nikon Stereo Microscope) equipped with a digital camera (Celestron 5MP CMOS Digital USB Microscope Imager). Each treatment consisted of one explant per tube, with 5–10 tubes per treatment. All viable, uncontaminated plantlets were retained for analysis. Tubes exhibiting microbial contamination or plantlet death due to handling damage were excluded from all measurements. To ensure consistency between replicate experiments, the physiological responses observed in MSO controls were compared by a Student’s t-test with no significant differences observed (p > 0.05).

### 2.4 Sample preparation for metabolite analysis

Culture samples were collected from gently washed roots and shoots following physiological observation. Shoots and roots were separated using a sterile razor blade, and each tissue type was transferred into a pre-weighed 1.5 mL microcentrifuge tube (Eppendorf). Fresh weight was recorded on an analytical balance (Explorer Pro, Ohaus), after which samples were immediately flash frozen in liquid nitrogen and stored at −80 °C until extraction.

Metabolite extraction was performed as previously described (Gaudet et al., 2026). Briefly, frozen tissues were ground directly in the microcentrifuge tubes using a disposable micro-pestle (Sigma-Aldrich) attached to a handheld vibration tool. Ground tissue was mixed at a 1:4 (w/v) ratio with 80% methanol containing 0.5 N trichloroacetic acid (TCA), vortexed for 1 min, and centrifuged at 5000 × g for 5 min. The resulting supernatant was transferred to 0.22 µm centrifugal filter units (Ultrafree-MC, PVDF membrane; Millipore) and centrifuged at 16,000 × g for 3 min. Filtered extracts were stored at 4 °C and analyzed within 48 hours.

### 2.5 UHPLC separation of kynurenine pathway metabolites

Extracts were transferred to autosampler vials (Canadian Life Sciences) and placed in the cooled autosampler (Vanquish Split Sampler, Thermo, at 8°C) until injection (10 µL). Kynurenine pathway metabolites were separated (Accucore Hypersil Gold pentafluorophenyl (PFP) column (150 × 3 mm, 2.6 µm particle size; Thermo Fisher Scientific) at 30°C with a constant flow rate of 0.5 mL min^−1^ delivered by a binary solvent system (Vanquish UHPLC, Thermo) consisting of (A) 0.1% (v/v) formic acid in water and (B) acetonitrile (UHPLC-MS grade, Thermo). Metabolite separation was previously optimized, and validation parameters were previously described (Gaudet et al., 2026). The standard gradient for separation of IAA, KYN and related metabolites is as follows: 0–2.0 min, 95% A / 5% B; 2.0–3.5 min, 70% A / 30% B; 3.5–6.5 min, 50% A / 50% B; 6.5–8.25 min, 5% A / 95% B; 8.25–10.5 min, 5% A / 95% B; 10.5–10.65 min, returned to 95% A / 5% B; 10.65–12.5 min, re-equilibration at 95% A / 5% B. The total run time was 12.5 min.

#### High-resolution mass spectrometric detection

Metabolites were detected and quantified using the previously validated methods (Gaudet et al., 2026). In brief, metabolites were ionized (heated electrospray ionization (HESI-II) source operated in positive ion mode), filtered and identified by high-resolution mass spectrometry (Q-Exactive Orbitrap mass spectrometer; Thermo). Full MS data were acquired at high resolution followed by data-dependent acquisition (DDA) scans for fragmentation confirmation. Parameters were optimized previously (Gaudet et al., 2026) as follows: spray voltage, 3.5 kV; capillary temperature, 269 °C; sheath gas flow rate, 53; auxiliary gas flow rate, 14; sweep gas flow rate, 3; auxiliary gas heater temperature, 438 °C; and S-lens RF level, 50. Full MS data were acquired at a resolution of 70,000 over a scan range of m/z 50–600, with data-dependent MS^2^ acquisition performed at a resolution of 17,500. Chromatographic peaks were extracted and integrated from the raw data files using TraceFinder (Thermo Fisher Scientific) and Skyline (MacCoss Lab Software). KYN, KYNA and 3-HAA were detected and quantified by previously published fully validated analytical methods using retention time, accurate mass, and DDA fragmentation matching to authentic standards (Table 1). Reference standards of KMO Inhibitor II (JM6), PF-04859989, and RO-61-8048 were identified by their characteristic retention times and m/z values to determine uptake in SJW but were not quantified by a validated analytical method (Table 1).

**Table 1:** Optimized parameters for high-resolution mass spectrometry detection and quantification of kynurenine pathway compounds*.

| Compound | Monoisotopic mass (m/z) | M + H (m/z) | Retention time in sample (min) | Lower Limit of Detection (pg on column) | Lower Limit of Quantification (pg on column) | %Recovery |
| --- | --- | --- | --- | --- | --- | --- |
| Kynurenine (KYN) | 208.08479 | 209.0921 | 4.05 | 10 | 25 | 78% |
| Kynurenic acid (KYNA) | 189.04559 | 190.0499 | 5.61 | 10 | 25 | 96% |
| 3-Hydroxyanthranilic acid (3-HAA) | 153.04259 | 154.0504 | 4.68 | 25 | 50 | 94% |
| KMO inhibitor II, JM6<sup>a</sup> | 518.1312 | 519.1385 | 7.9 | NA | NA | NA |
| PF-04859989<sup>a</sup> | 178.0745 | 179.0818 | 4.8 | NA | NA | NA |
| RO-61-8048<sup>a</sup> | 421.0390 | 422.0463 | 8.41 | NA | NA | NA |

\* Limit of detection and limit of quantification were determined by repeated injection of a dilution series of eight concentrations spanning 10 – 10,000 pg on column representing the linear range for major metabolites.
<sup>a</sup> Limit of detection and quantification were not established for pharmaceutical inhibitors as this data was used to confirm presence.

### 2.6 Experimental design and statistical analysis

Each treatment consisted of one explant per culture tube, with 5–10 tubes per treatment. A total of four independent experimental runs were conducted. All viable, uncontaminated plantlets were retained for analysis while tubes exhibiting contamination or plantlet death were excluded. Rooting frequency, maximum root length, shoot height, and metabolite concentrations were analyzed per plantlet, with three biological replicates used for chemical analysis in each treatment.

All statistical analyses were performed in R (version 4.4.0; R Core Team) using base R functions and the *emmeans* and *multcomp* packages (Hothorn et al., 2008; Searle et al., 1980). Rooting frequency (binary) was analyzed using a generalized linear model with binomial error distribution. Treatment effects were evaluated using Tukey-adjusted contrasts for pairwise comparisons among treatments. Physiological measurements (root length and shoot height) were evaluated using Dunnett contrasts comparing each treatment with the MSO control, whereas metabolite concentrations among multiple treatments were compared using Tukey’s HSD test for pairwise comparisons. Comparisons involving only two groups (e.g., root versus shoot metabolite concentrations under MSO conditions) were analyzed using a two-sample t-test. Pathway diagrams were prepared in Inkscape (Inkscape Project, https://inkscape.org), and chemical structures were drawn using ACD/ChemSketch (Advanced Chemistry Development, Inc. (ACD/Labs), Toronto, ON, Canada). Statistical significance was defined as α = 0.05 for all analyses.

## 3. Results

### 3.1 Establishment of the Bioassay System

Preliminary experiments established PGR rates for application. *De novo* root regeneration was observed on SJW plantlets grown on basal media in the absence of any exogenous PGRs. SJW plantlets grown on media supplemented with 10 μM IAA form a callus ball at the cut surface (Fig. 2A). Supplementation of IAA in the culture media induced a stunted morphology with shorter shoots, multiple shoots and smaller leaves. The morphology of plants grown on media supplemented with 10 μM KYN was visually consistent with the basal media control (Fig. 2A). Supplementation of the IAA media with KYN metabolism inhibitors reversed the growth morphology observed with IAA alone (Fig. 2A). Rooting frequency was significantly affected by treatment (Fig. 2B). MSO control explants exhibited a rooting frequency of 0.63 ± 0.07. Treatment with IAA significantly reduced rooting frequency to 0.09 ± 0.07, whereas KYN (0.83 ± 0.06), JM6 (0.88 ± 0.08), PF (0.57 ± 0.10), RO (0.69 ± 0.09), and the combined IAA treatments (IAA/JM6 0.27 ± 0.11, IAA/PF 0.25 ± 0.11, IAA/RO 0.40 ± 0.13) did not differ significantly from MSO controls at the p ≤ 0.05 level. Supplementation with KYN or inhibition of KYN metabolism did not significantly affect root growth. IAA inhibition of root growth was partially recovered by inhibition of KYN metabolism but not significantly different (Fig. 2B). Internodal length differed among treatments (Fig. 2C). MSO explants exhibited an internodal length of 0.400 ± 0.180 cm per node. KYN significantly increased internodal length to 0.552 ± 0.198 cm per node, and IAA/PF also produced a significant increase (0.566 ± 0.278 cm per node). Other treatments did not differ significantly from MSO control.

**Figure 2.**
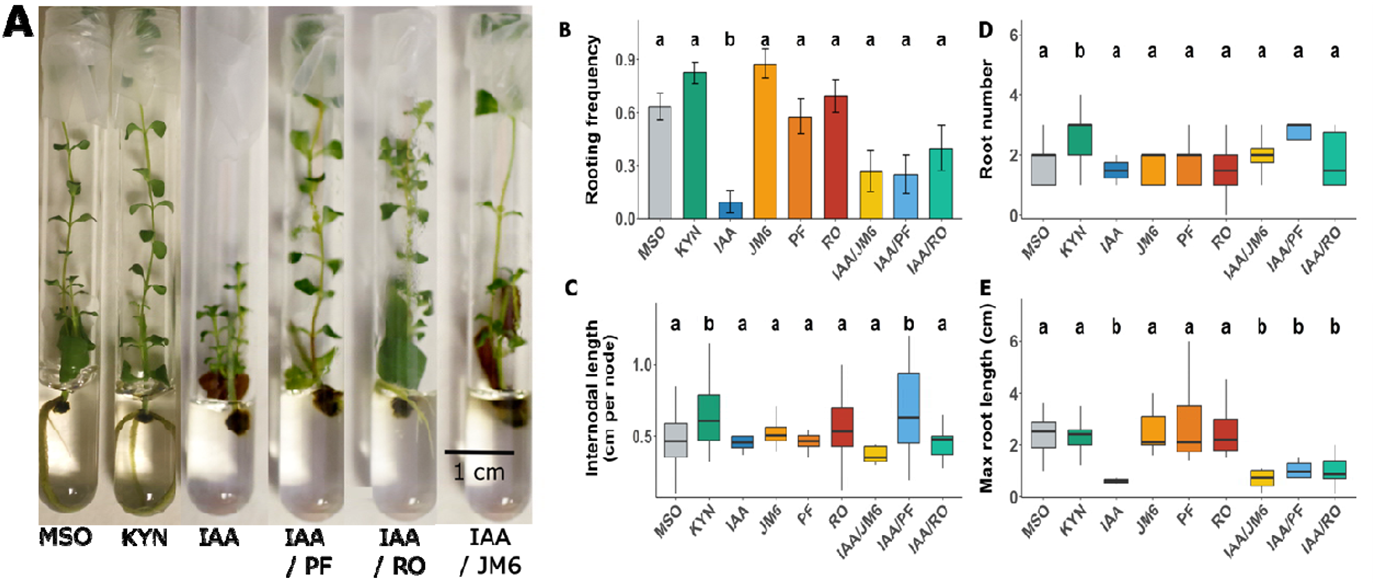
Effects of kynurenine pathway metabolites and inhibitors on *in vitro* root and shoot development in *Hypericum perforatum*. (A) Representative images of explants cultured on MSO (control), kynurenine (KYN), indole-3-acetic acid (IAA), and IAA combined with inhibitors (IAA + JM6, IAA + PF-04859989 [PF], and IAA + RO 61-8048 [RO]). Scale bar = 1 cm (B) Rooting frequency, (C) internodal length (cm per node), (D) root number, and (E) maximum root length (cm) of explants under each treatment. For rooting frequency (B), bars represent mean proportion rooted ± SE. For (C–E), boxplots represent median (center line), interquartile range (box), and range (whiskers). Differences relative to the MSO control were evaluated using Dunnett-adjusted contrasts (p < 0.05; n = 12–18 per treatment).

Root number differed among treatments (Fig. 2D). MSO explants produced 1.45 ± 0.97 roots on average. KYN significantly increased root number to 2.52 ± 0.94 roots, whereas all other treatments did not differ significantly from MSO control. Maximum root length differed among treatments (Fig. 2E). IAA significantly reduced maximum root length relative to MSO, and similar reductions were observed for the combined treatments IAA/JM6, IAA/PF, and IAA/RO. Treatments with KYN or inhibitors alone did not differ significantly from MSO control.

### 3.1 Kynurenine pathway metabolites are present in roots and shoots of *Hypericum perforatum*

Assays of explants grown on MSO media determined that the KYN pathway metabolites KYN, KYNA, and 3-HAA were detected in both root and shoot tissues of *H. perforatum* (Fig. 3A–C). KYN, KYNA and 3-HAA were quantified at significantly higher concentrations in shoots than in roots (Fig. 3). In shoots, KYNA was quantified at 247 ± 110 ng g^−1^ FW (Fig. 3A), KYN was quantified at 234 ± 52 ng g^−1^ FW (Fig. 3B), and 3-HAA was quantified at 462 ± 135 ng g^−1^ FW (Fig. 3C). In root tissues, KYNA, KYN and 3-HAA were quantified at 41 ± 4 ng g^−1^ FW (Fig. 3A), 98 ± 19 ng g^−1^ FW (Fig 3B), and 133 ± 38 ng g^−1^ FW (Fig. 3C) respectively.

**Figure 3.**
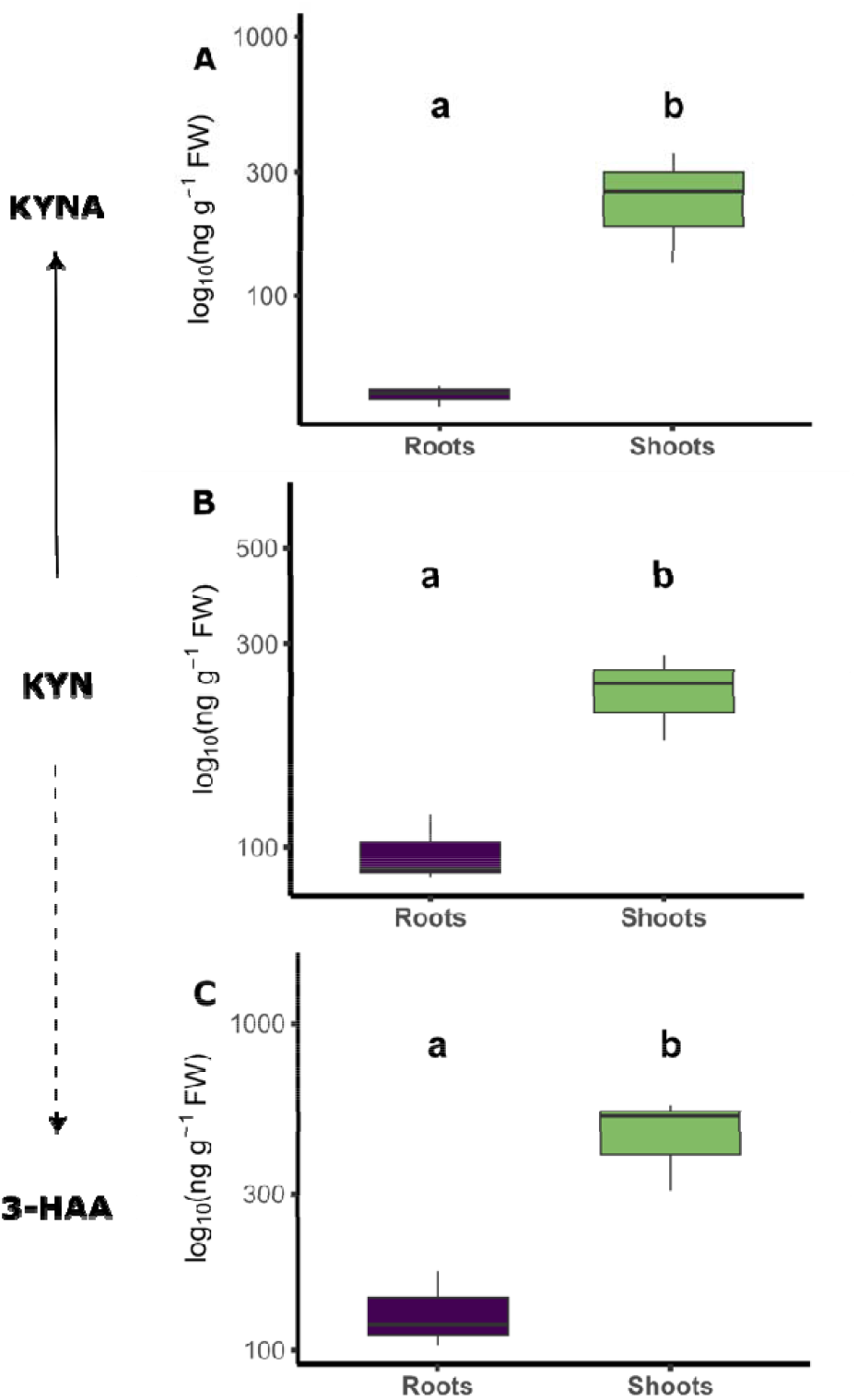
Kynurenine pathway metabolites in roots and shoots of *Hypericum perforatum* plantlets grown on MSO medium. (A) Kynurenic acid (KYNA), (B) kynurenine (KYN), and (C) 3-hydroxyanthranilic acid (3-HAA). Concentrations are expressed as ng g^−1^ fresh weight (FW). Boxplots represent median (center line), interquartile range (box), and range (whiskers). Asterisks indicate significant differences between roots and shoots for each metabolite (t-test, p < 0.05; n = 3 per tissue). The simplified pathway schematic indicates established conversions (solid lines) and putative or unresolved steps (dotted lines).

### 3.2 Kynurenine pathway inhibitors JM6 and RO increase KYN accumulation in tissues

To confirm that inhibitors could be taken up by plant tissues, we first confirmed their presence in tissues through UHPLC-MS analysis. Inhibitors PF-04859989, RO-61-8048, and KMO inhibitor

II, JM6 were detected in SJW explants following treatment (Fig. 4A-C). The relative signal intensity was significantly higher in roots than in shoots for each of the inhibitors indicating both uptake and transport in the explants (Fig. 4D-F).

**Figure 4.**
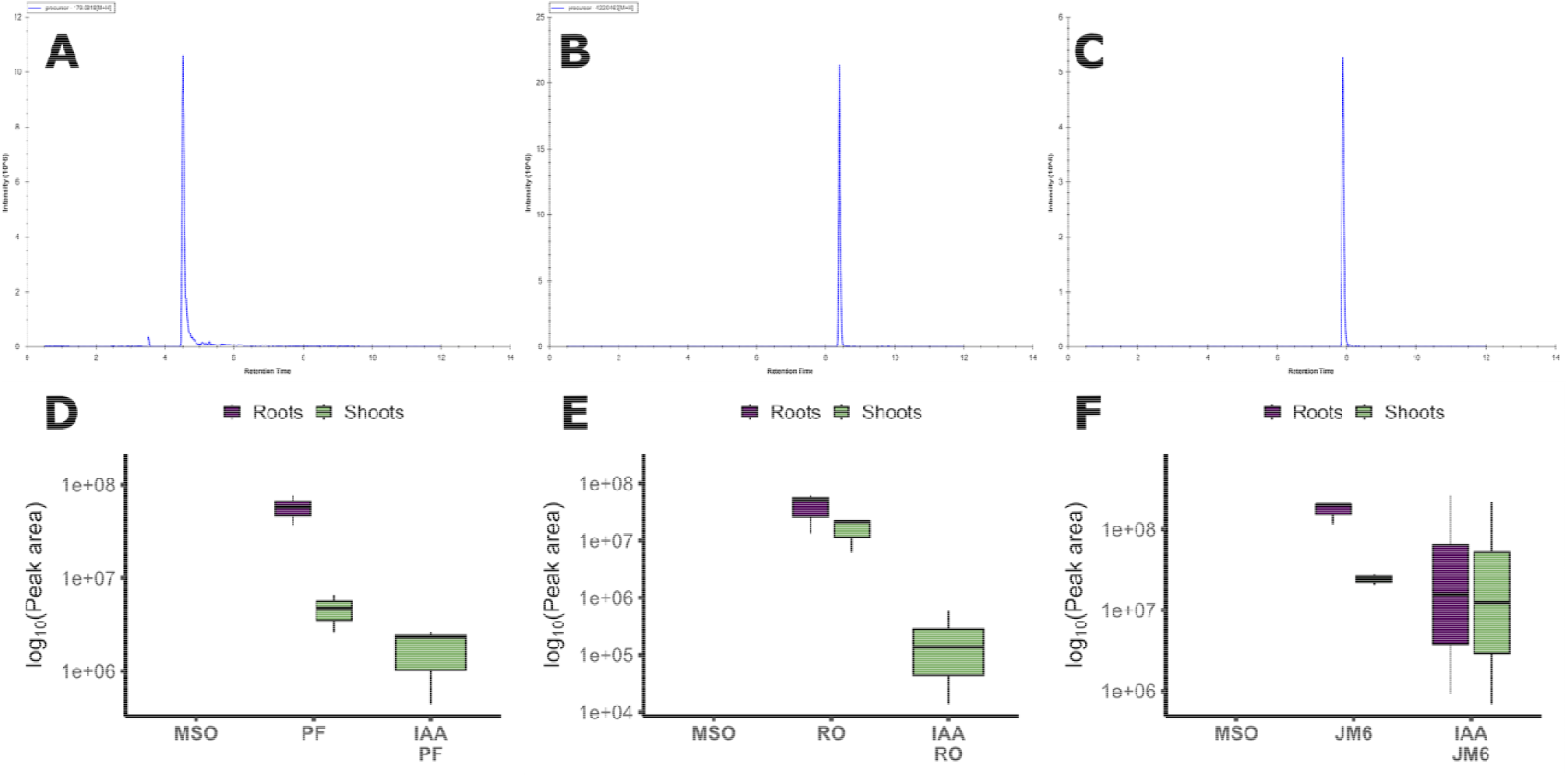
Detection and relative abundance of kynurenine pathway inhibitors in *Hypericum perforatum* tissues. (A–C) Representative extracted ion chromatograms (EICs) of PF-04859989 (A), RO 61-8048 (B), and JM6 (KMO inhibitor II) (C) detected in plant tissue by LC–HRMS. Each panel shows the precursor ion trace at the expected m/z and retention time. (D–F) Relative abundance of PF (D), RO (E), and JM6 (F) in roots and shoots following treatment with MSO (control), inhibitor alone, or IAA + inhibitor. Peak areas are shown as log□□-transformed values. Boxplots represent median (center line), interquartile range (box), and range (whiskers). Signals corresponding to each inhibitor were observed in treated tissues and were not detected in MSO controls. Detection was also observed in IAA co-application treatments.

### 3.3 Exogenous KYN increases KYNA in roots, but does not affect 3-HAA

#### 3.3.1 De novo shoot organogenesis

KYNA concentrations did not differ significantly between KYN-treated shoots (890 ± 332 ng g^−1^ FW) and MSO controls (247 ± 110 ng g^−1^ FW) at the p ≤ 0.05 level due to relatively high variability in the measures (Fig. 5A). KYN concentration was significantly higher in shoots of KYN-treated explants (703 ± 266 ng g^−1^ FW) as compared with MSO controls (234 ± 52 ng g^−1^ FW) (Fig. 5B). Similarly, 3-HAA concentrations did not significantly differ between KYN-treated shoots (640 ± 161 ng g^−1^ FW) and MSO controls (462 ± 135 ng g^−1^ FW) (Fig. 5C).

**Figure 5.**
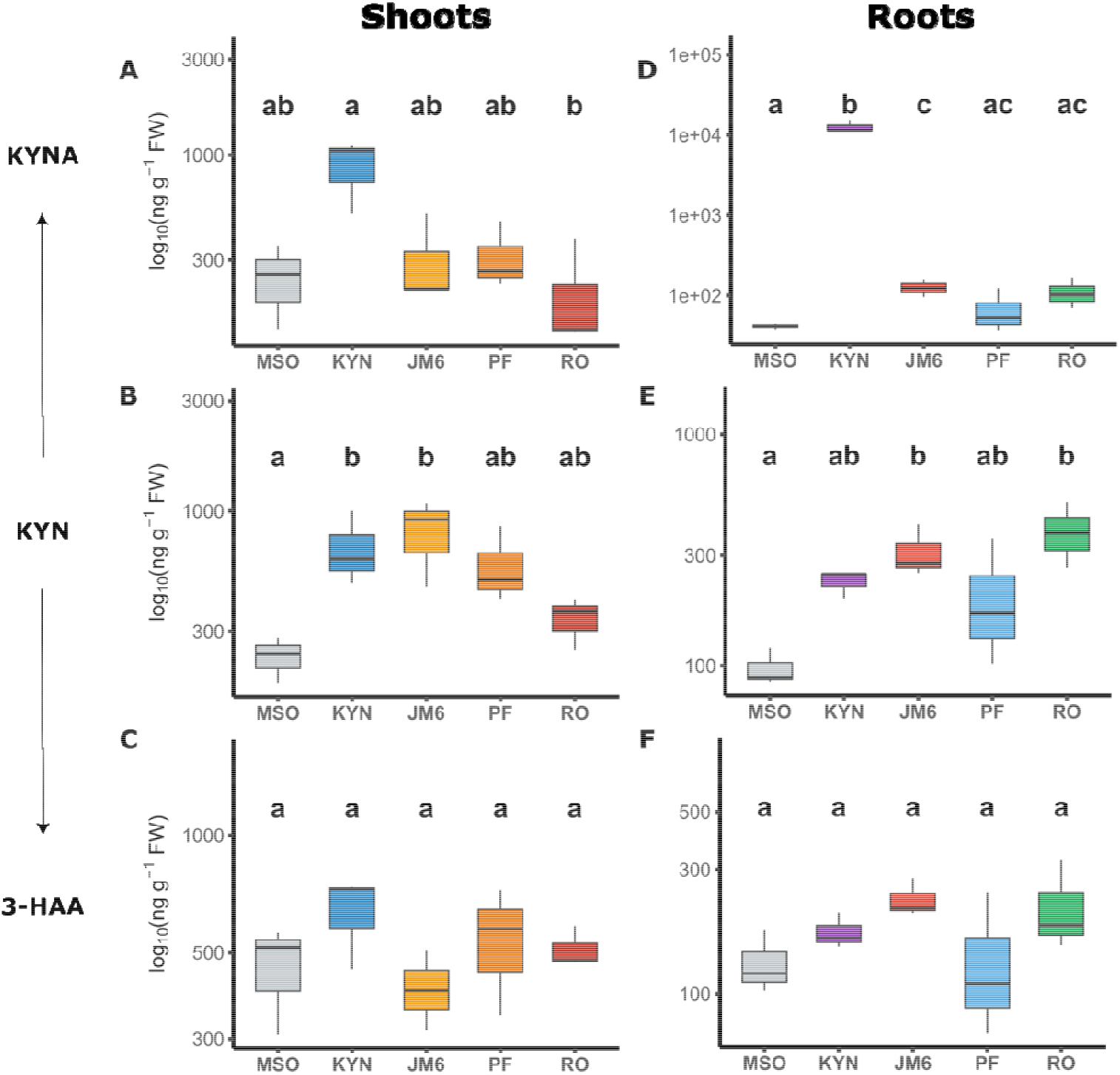
Concentration of kynurenine pathway metabolites in roots and shoots following treatment with MSO, kynurenine (KYN), KMO inhibitor II (JM6), PF-04859989 (PF), and RO 61-8048 (RO). (A–C) Shoots and (D–F) roots. (A, D) Kynurenic acid (KYNA), (B, E) kynurenine (KYN), and (C, F) 3-hydroxyanthranilic acid (3-HAA). Concentrations are shown as log□□ (ng g^−1^ FW). Boxplots represent median (center line), interquartile range (box), and range (whiskers). Different letters indicate significant differences among treatments within each tissue and metabolite (one-way ANOVA followed by Tukey’s HSD, p < 0.05; n = 3).

#### 3.3.2 De novo root organogenesis

KYNA concentrations were significantly higher in roots of KYN-treated explants (12,706 ± 2,338 ng g^−1^ FW) as compared to the roots of the MSO control explants (41 ± 4 ng g^−1^ FW) (Fig. 5D) but the KYN concentrations were not significantly different between roots of MSO control explants (98 ± 19 ng g^−1^ FW) and the KYN-treated explants (231 ± 32 ng g^−1^ FW) (Fig. 4E). KYN treatment did not significantly affect 3-HAA concentrations in explant roots (174 ± 28 ng g^−1^ FW) as compared with control roots (133 ± 38 ng g^−1^ FW) (Fig. 5F).

#### 3.3.3 Effects of pathway inhibitors

Pharmaceutical drug prototypes designed to inhibit human KYN metabolism significantly impacted plant KYN metabolism. The high-affinity inhibitor of KMO in humans, RO-61-8048 did not significantly affect KYNA or 3-HAA levels in plants (Fig. 5). RO-61-8048 significantly increased the concentration of KYN in roots (383 ± 123 ng g^−1^ FW) as compared to MSO control explants controls (98 ± 19 ng g^−1^ FW) (Fig. 5E). The stabilized version of RO-16-8048, JM6, has the same enzyme target in mammals but induced different responses in planta. SJW explants had significantly higher KYN levels in both shoots (823 ± 315 ng g^−1^ FW) and roots (313 ± 84 ng g^−1^ FW) in response to JM6 (Fig. 5B&E). In roots, KYNA concentrations were also significantly increased by JM6 exposure (127 ± 32 ng g^−1^ FW) as compared with the MSO control explants (41 ± 4 ng g^−1^ FW) (Fig. 5D). PF-04859989 was selected as a potential high-affinity inhibitor of human KAT but was a less effective mediator of KYN metabolism in planta (Fig. 5).

### 3.4 Exogenous IAA increases KYN and 3-HAA in shoots

We investigated the potential interactions of IAA and KYN pathways with combined treatments of supplementation and inhibition (Fig. 2 & 6). IAA significantly reduced rooting frequency in the bioassay but addition of the inhibitors PF-04859989, RO-61-8048 and JM6 partially recovered the de novo rooting response (Fig. 2A). Quantification of KYNA did not reveal significant differences in shoots in response to the combined auxin + inhibitor treatments (Fig. 6A), with values of 247 ± 110 ng g^−1^ FW in MSO controls, 529 ± 119 ng g^−1^ FW in IAA-treated tissues, 521 ± 227 ng g^−1^ FW in IAA/JM6-treated tissues, 345 ± 65 ng g^−1^ FW in IAA/PF-treated tissues, and 562 ± 387 ng g^−1^ FW in IAA/RO-treated tissues. KYN concentrations were significantly different in shoots as a result of IAA + inhibitor exposures (Fig. 5B). MSO control explants had 234 ± 52 ng g^−1^ FW of KYN, while IAA-treated tissues accumulated 656 ± 82 ng g^−1^ FW and IAA/JM6-treated tissues accumulated 1279 ± 201 ng g^−1^ FW. IAA/PF (641 ± 390 ng g^−1^ FW) and IAA/RO (588 ± 246 ng g^−1^ FW) (Fig. 6B). Shoots of explants grown on MSO media contained 462 ± 135 ng g^−1^ FW 3-HAA (Fig. 6C). Combinations of IAA and inhibitors significantly increased 3-HAA in shoots of all other treatments (IAA - 1298 ± 195 ng g^−1^ FW; IAA+JM6 - 967 ± 121 ng g^−1^ FW; IAA + PF - 1027 ± 27 ng g^−1^ FW; and IAA + RO - 1000 ± 256 ng g^−1^ FW (Fig. 6C)

**Figure 6.**
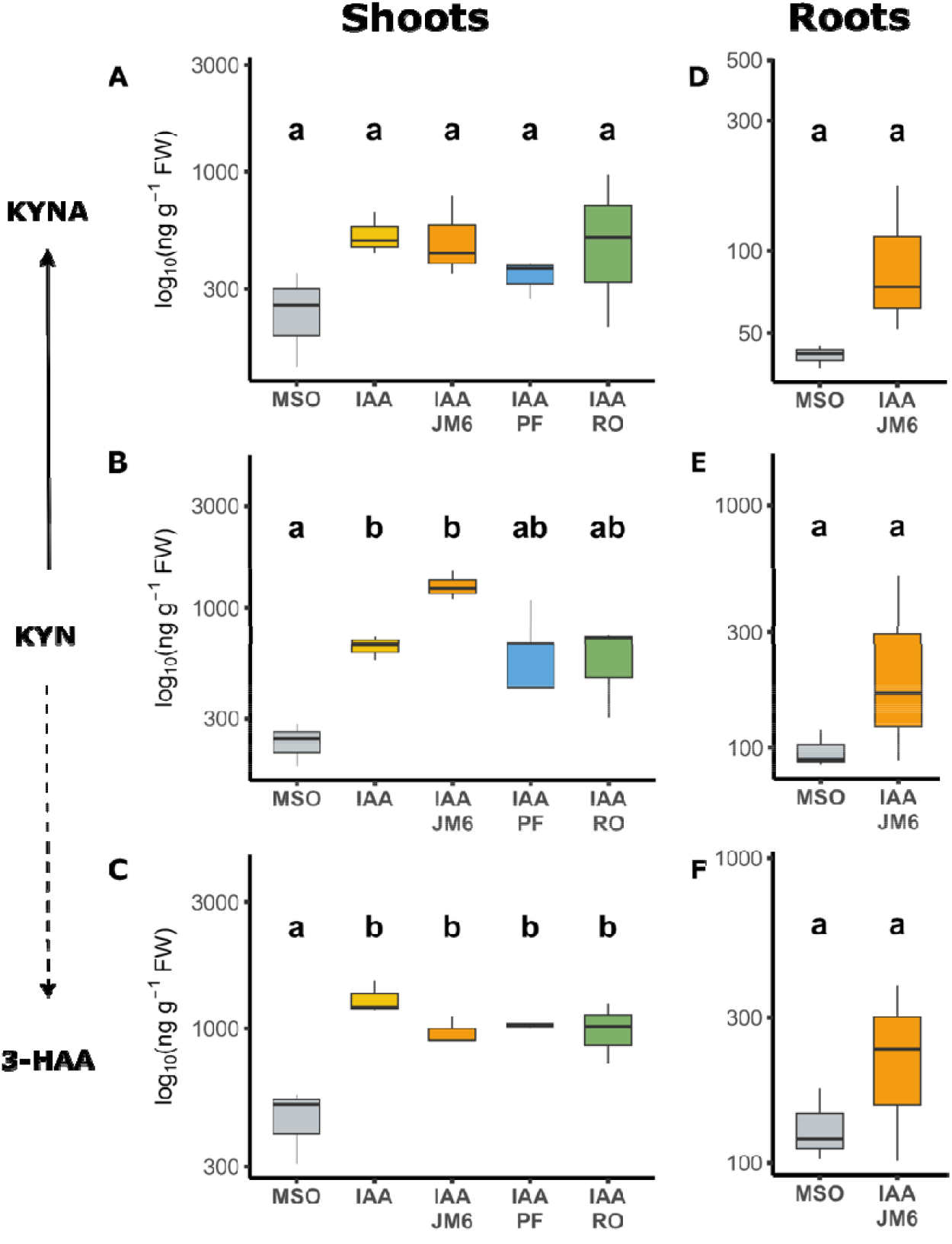
Effects of indole-3-acetic acid (IAA) and kynurenine pathway inhibitors on kynurenine pathway metabolite concentrations in *Hypericum perforatum*. Concentrations of (A, D) kynurenic acid (KYNA), (B, E) kynurenine (KYN), and (C, F) 3-hydroxyanthranilic acid (3-HAA) in shoots (A–C) and roots (D–F) of explants cultured on MSO (control), IAA, or IAA combined with kynurenine pathway inhibitors (IAA + JM6, IAA + PF-04859989, and IAA + RO 61-8048). Concentrations are shown as log□□ (ng g^−1^ FW). Boxplots represent median (center line), interquartile range (box), and range (whiskers). For shoots (A–C), different letters indicate significant differences among treatments (one-way ANOVA followed by Tukey’s HSD, p < 0.05; n = 3). For roots (D–F), differences relative to the MSO control were evaluated using Dunnett-adjusted contrasts (p < 0.05; n = 3).

In roots, metabolites were detected in explants grown on MSO and IAA + JM6 media due to limited root growth and lack of tissue harvested from other treatments (Fig. 2D–F). While data appear to show a trend toward higher amounts of KYNA, KYN and 3-HAA in IAA + JM6 treated explants, these data were not significant at p ≤ 0.05 (Fig. 6D, E, F). KYNA was quantified in roots grown on MSO (41 ± 4 ng g^−1^ FW) and IAA/JM6-treated explants (100 ± 65 ng g^−1^ FW) (Fig. 6D). KYN was also quantified in roots of explants grown on MSO media (98 ± 19 ng g^−1^ FW) and IAA/JM6-treated explants (258 ± 228 ng g^−1^ FW) (Fig. 6E). 3-HAA was quantified in roots of explants grown on MSO (133 ± 38 ng g^−1^ FW) and IAA/JM6-treated tissues (240 ± 140 ng g^−1^ FW) (Fig. 6F).

## 4. Discussion

We hypothesized that KYN metabolism mediates organogenesis in St. John’s wort. To test this hypothesis, we established a bioassay system for investigations of KYN pathway activity and interactions with auxin metabolism. SJW bioassays have previously been used for studies of auxin-induced morphogenesis (Murch et al., 2001; Murch and Saxena, 2006) and the indoleamine plant growth regulators serotonin and melatonin (Erland et al., 2018, 2019). Plant regeneration is regulated by the relative ratios of auxins and cytokinins (Skoog and Miller, 1957), excision, isolation and stress, and cross-talk among growth-regulating signals (Zhao, 2010). Plant cell strategies for maintaining auxin homeostasis include polar auxin transport, conjugation, deactivation, sequestration and catabolism (Guo et al., 2022; Maruyama and Ikeuchi, 2026). The modern understanding of plant growth regulation is a complex network of cross-talk between small signalling molecules, with auxin at the core of this network (Maruyama and Ikeuchi, 2026).

The most significant finding of the current work is the discovery of a potential role of KYN in auxin catabolism and plant growth. Externally applied KYN accumulated in roots and was transported to shoots. Our data show that KYN is metabolized to KYNA and 3-HAA. This pattern may reflect KYNA accumulation as a terminal product, while 3-HAA is further metabolized toward NAD□ biosynthesis (Savitz, 2020). Kynurenine occupies a central branch point in the pathway and is metabolized either toward kynurenic acid via KAT or toward downstream oxidative metabolites via KMO (Badawy, 2017; Stone et al., 2013). To determine whether this branch structure explains the observed metabolite distribution, we applied mammalian pharmacological inhibitors targeting KAT and KMO (Jacobs et al., 2017). We confirmed that KMO Inhibitor II (JM6), RO-61-8048, and PF-04859989 were absorbed from the media and affected plant morphogenesis, KYN metabolism and auxin catabolism. The contrasting effects of KYN and IAA on rooting and internodal growth further support the conclusion that KYN metabolism participates in organogenic regulation.

Previous studies have shown that KYN may control auxin biosynthesis by inhibiting TAA1 (He et al., 2011; Maruyama and Ikeuchi, 2026). Anthranilic acid, an early precursor of IAA, has also been shown to regulate root gravitropism through effects on PIN polarity and auxin distribution independently of changes in IAA levels (Doyle et al., 2019). Auxin catabolism has evolved in at least two distinct evolutionary lineages in microbes (Conway et al., 2022), but plant catabolic mechanisms remain unelucidated. Chemical oxidation of tryptophan produces KYN and related metabolites in the presence of iron (FeCl□) and hexanal, and the degradation can be mediated by plant polyphenols and antioxidants (Salminen et al., 2008). Our data provide evidence that auxin catabolism may contribute to KYN-associated metabolite accumulation, thereby providing feedback inhibition on TAA1 to control auxin homeostasis. Together, these findings support an auxin homeostasis model in which IAA metabolism, oxidative transformation and KYN pathway activity interacts within the tryptophan-derived metabolic network of SJW (Fig. 7). Further research is required to fully elucidate the interactions between KYN and auxin in plant growth.

**Figure 7.**
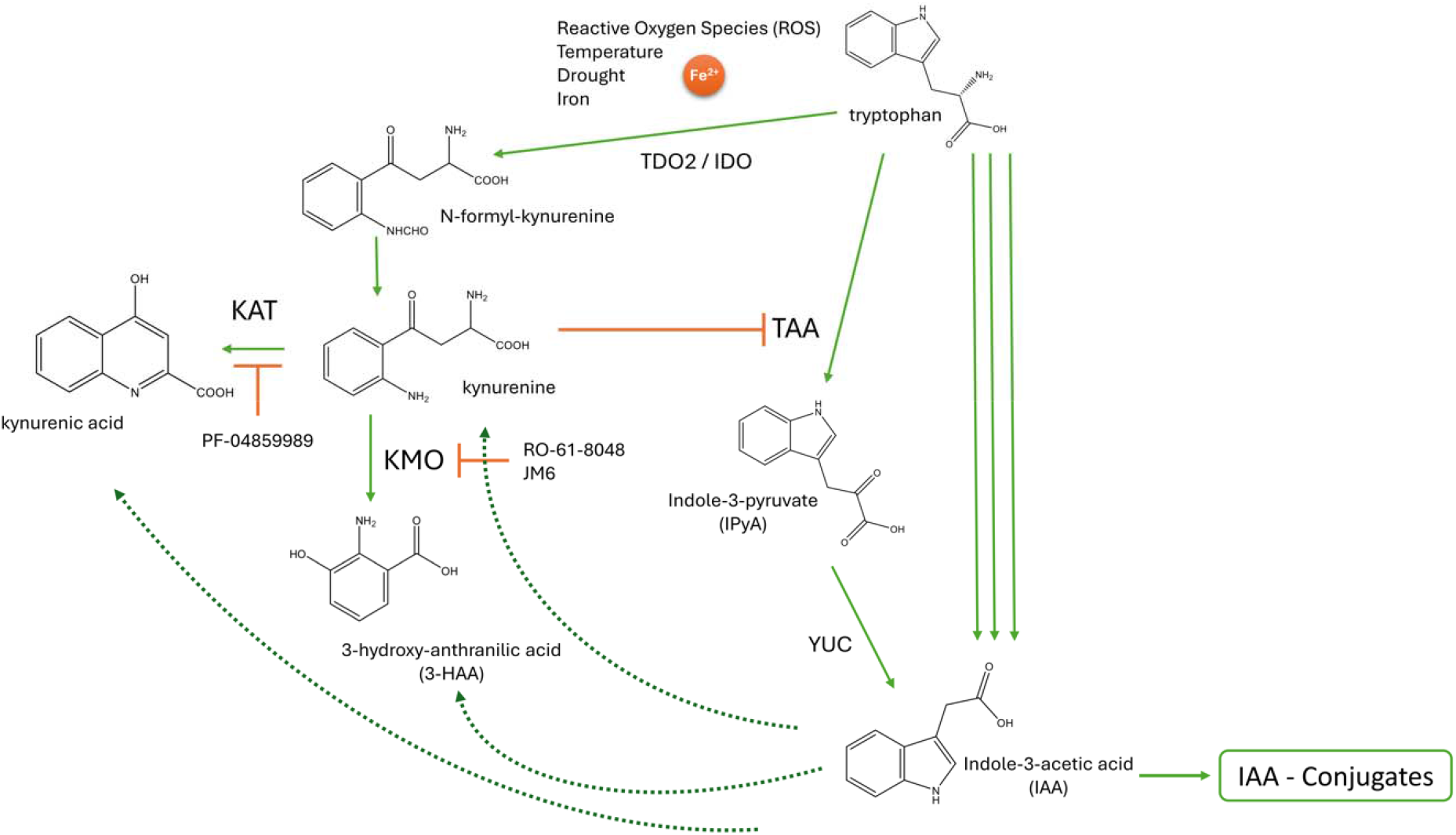
Proposed auxin homeostasis model linking IAA metabolism with kynurenine-associated tryptophan metabolism in *Hypericum perforatum*. Indole-3-acetic acid (IAA) is primarily synthesized from tryptophan through the indole-3-pyruvate (IPyA) pathway via tryptophan aminotransferase (TAA) and YUCCA flavin monooxygenase (YUC). Free IAA may be regulated through conjugation, catabolism, oxidative transformation and through feedback effects on tryptophan-derived metabolism. Kynurenine pathway metabolism proceeds through N-formyl-kynurenine and kynurenine, which occupies a central branch point between kynurenic acid formation via kynurenine aminotransferase (KAT) and downstream oxidative metabolism toward 3-hydroxyanthranilic acid (3-HAA) via kynurenine monooxygenase (KMO). Reactive oxygen species (ROS), temperature, drought, iron, and Fe^2+^ are shown as potential stress and redox inputs that may influence auxin and kynurenine-associated metabolism. The pharmacological inhibitors used in this study are shown at their proposed targets: PF-04859989 at KAT, and RO-61-8048 and JM6 at kynurenine monooxygenase (KMO). Dashed arrows indicate proposed interactions linking auxin catabolism or oxidative transformation with kynurenine-associated metabolite accumulation and potential feedback on tryptophan-dependent auxin biosynthesis.

## Acknowledgements

The authors acknowledge funding support from the Natural Sciences and Engineering Council (NSERC) Canada (04928) and thank Supra Research and Development for technical support related to UHPLC-MS analysis.

## CRediT Author Statement

**Daniel Gaudet:** Conceptualization, Methodology, Investigation, Formal analysis, Data curation, Visualization, Writing. **Alisha Greene:** Methodology, Investigation **Susan J. Murch:** Conceptualization, Methodology, Supervision, Resources, Writing **Lauren A.E. Erland:** Conceptualization, Methodology, Project administration, Supervision, Resources, Writing

