## Supplementary Data for "Auxin is metabolized through kynurenine in *Hypericum perforatum* L"

**Supplemental Table S1:** Detection of kynurenine pathway metabolites (kynurenine (KYN), kynurenic acid (KYNA) and 3-hydroxyanthranilic acid (3-HAA)) in plant species reported in previous literature.

| Species | KP metabolites detected | Reference(s) |
|---|---|---|
| *Hypericum perforatum* | KYN, KYNA, 3-HAA | Gaudet et al. (2026) |
| *Cannabis sativa* | KYN, KYNA | Russo et al. (2022) |
| *Taraxacum officinale* | KYNA | Turski et al. (2011) |
| *Urtica dioica* | KYNA | Turski et al. (2011) |
| *Chelidonium majus* | KYNA | Turski et al. (2011) |
| *Aesculus hippocastanum* | KYNA | Turski et al. (2011) |
| *Matricaria chamomilla* | KYNA | Turski et al. (2011) |
| *Linum usitatissimum* | KYNA | Turski et al. (2011) |
| *Crataegus oxyacantha* | KYNA | Turski et al. (2011) |
| *Viola tricolor* | KYNA | Turski et al. (2011) |
| *Melissa officinalis* | KYNA | Turski et al. (2011) |
| *Tilia cordata* | KYNA | Turski et al. (2011) |
| *Brassica oleracea* var. *botrytis* | KYNA | Turski et al. (2009, 2011) |
| *Brassica oleracea* var. *italica* | KYNA | Turski et al. (2009, 2011) |
| *Solanum tuberosum* | KYNA | Turski et al. (2009); Turska et al. (2022 review) |
| *Daucus carota* | KYNA | Turski et al. (2009) |
| *Capsicum annuum* | KYNA | Turski et al. (2009) |
| *Malus domestica* | KYNA | Turski et al. (2009); Turska et al. (2022 review) |
| *Cucumis sativus* | KYNA | Turski et al. (2009) |
| *Solanum lycopersicum* | KYNA | Turski et al. (2009); Turska et al. (2022 review) |
| *Pisum sativum* | KYNA | Turski et al. (2009) |
| *Allium cepa* | KYNA | Turski et al. (2009) |
| *Allium sativum* | KYNA | Turski et al. (2009) |
| *Zea mays* | KYNA | Turski et al. (2009) |
| *Hordeum vulgare* | KYNA | Turski et al. (2009) |
| *Oryza sativa* | KYNA | Turski et al. (2009) |
| *Brassica napus* | KYNA | Turski et al. (2009) |
| *Helianthus annuus* | KYNA | Turski et al. (2009) |
| *Ocimum basilicum* | KYNA | Turska et al. (2022 review) |
| *Thymus vulgaris* | KYNA | Turska et al. (2022 review) |
| *Castanea sativa* | KYNA | Turska et al. (2022 review) |

**Supplemental Table S2:** Optimized Parameters for High Resolution (OrbiTrap) Mass Spectrometry detection and quantification of kynurenine, kynurenic acid and 3-hydroxyanthrilic acid (as per Guadet et al., 2026).

Instrument Parameters:

(a) Mode: Positive
(b) Calibrators: Pierce LTQ Velos ESI Positive Ion Calibration Solution. Calibration was performed prior to analysis.
(c) Sheath gas flow rate: 53
(d) Aux gas flow rate: 14
(e) Sweep gas flow rate: 3
(f) Sheath, aux and sweep gas: Peak scientific Genius XE nitrogen generator
(g) Spray voltage: 3.5 kV
(h) Capillary temperature: 269 °C
(i) S lens Rf level: 50.0
(j) Aux gas heater temperature: 438 °C
(k) Divert valve: Eluent sent to the mass spectrometer from 0.5 to 12 mins. Sent to waste at other times.
(l) Acquisition mode: Full MS with ddMS2.
(m) Properties of full MS/ dd-MS2:
(n) Collision gas: Ultra-high purity nitrogen.

Detection Settings:

| | | |
|---|---|---|
| General | Runtime | 0.5 to 12 mins |
| | Polarity | Positive |
| | In-source CID | 0.0 eV |
| | Default charge state | 1 |
| Full MS | Microscans | 1 |
| | Resolution | 70,000 |
| | AGC target | 3e6 |
| | Maximum IT | 100 ms |
| | Number of scan ranges | 1 |
| | Scan range | 50 to 600 m/z |
| | Spectrum data type | Profile |
| dd-MS2/ dd-SIM | Microscans | 1 |
| | Resolution | 17,500 |
| | AGC target | 5e4 |
| | Maximum IT | 50 ms |
| | Loop count | 5 |
| | MSX count | 1 |
| | TopN | 5 |
| | Isolation window | 2.0 m/z |
| | Isolation offset | 0.0 m/z |

| | | |
|---|---|---|
| | (N)CE/ stepped (N) CE | nce: 17.5, 30, 42.5 |
| | Spectrum data type | profile |
| dd Settings | Minimum AGC target | 8e3 |
| | Intensity threshold | 1.6e5 |

**Supplementary Table S3: Preparation of the dilution series of analytical standards**

| Standard | Concentration (ng/mL) | Final Working Concentration (ng / mL) | Picograms on column |
|---|---|---|---|
| **0** | 10000000 | - | - |
| **1** | 1000000 | - | - |
| **1.5** | 500000 | - | - |
| **2** | 100000 | - | - |
| **2.5** | 50000 | - | - |
| **3** | 10000 | 1000 | 10,000 |
| **3.5** | 5000 | 500 | 5,000 |
| **4** | 1000 | 100 | 1,000 |
| **4.5** | 500 | 50 | 500 |
| **5** | 100 | 10 | 100 |
| **6** | 50 | 5 | 50 |
| **7** | 25 | 2.5 | 25 |
| **8** | 10 | 1 | 10 |

**Figure S1. Representative LC–HRMS chromatograms of kynurenine pathway inhibitor standards.**

Extracted ion chromatograms of PF-04859989 (A), RO 61-8048 (B), and KMO inhibitor II, JM6 (C) acquired under the same chromatographic and mass spectrometric conditions used for plant tissue analysis. Each inhibitor produced a distinct peak at its characteristic retention time. Signals were obtained from methanol standards at 50 ng $mL^{-1}$, confirming chromatographic separation and method sensitivity.

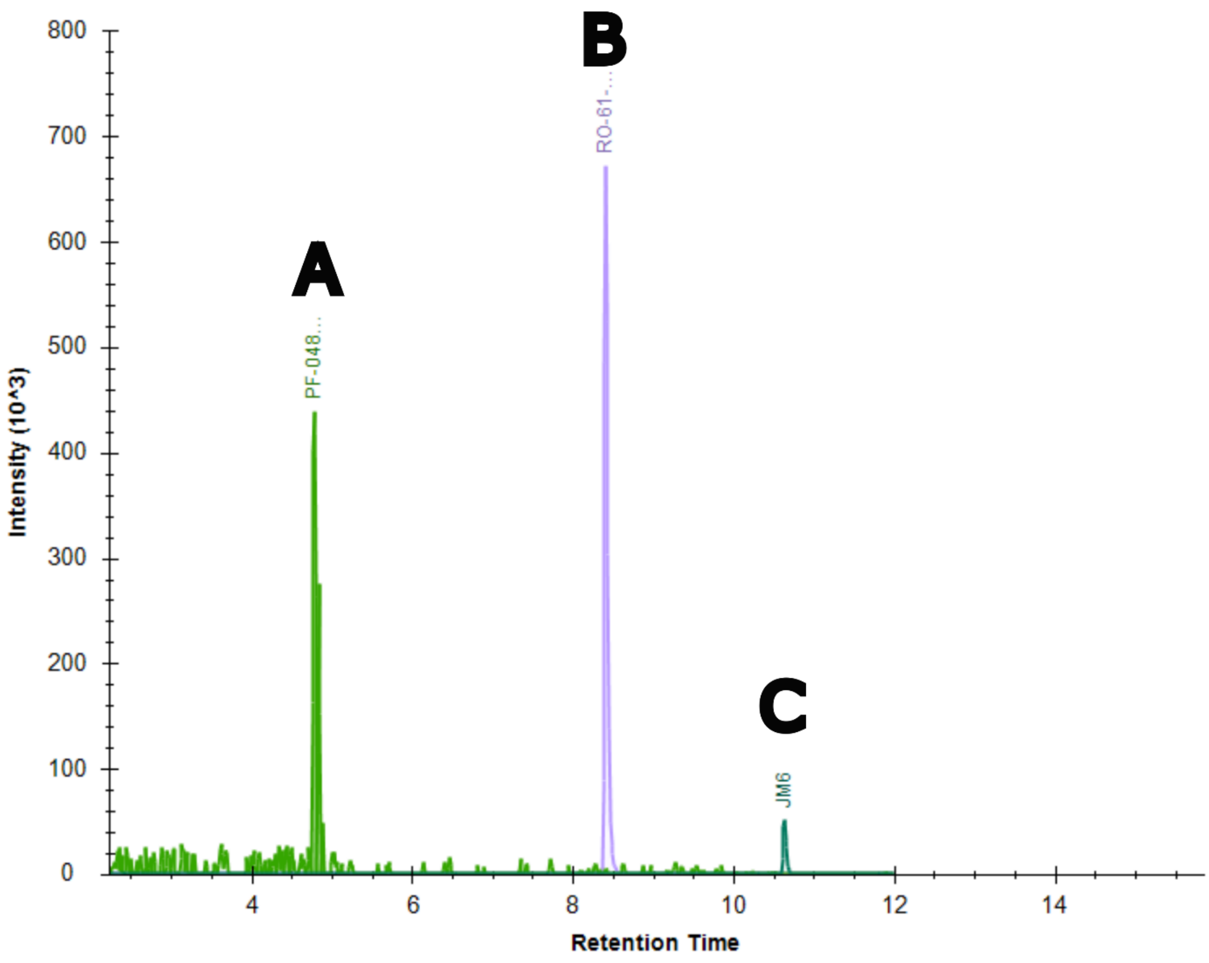

**Figure S2. Shoot developmental parameters under kynurenine pathway metabolite and inhibitor treatments.**

Number of internodes, (B) cumulative shoot height (cm), and (C) maximum shoot height (cm) of explants grown under MSO (control), KYN, IAA, inhibitor-only treatments (JM6, PF-04859989, RO 61-8048), and combined treatments with IAA (IAA + JM6, IAA + PF-04859989, IAA + RO 61-8048). Data are presented as boxplots showing median, interquartile range, and range. No treatments differed significantly from MSO based on Dunnett-adjusted comparisons ($p \geq 0.05$; n = 12–18 per treatment).

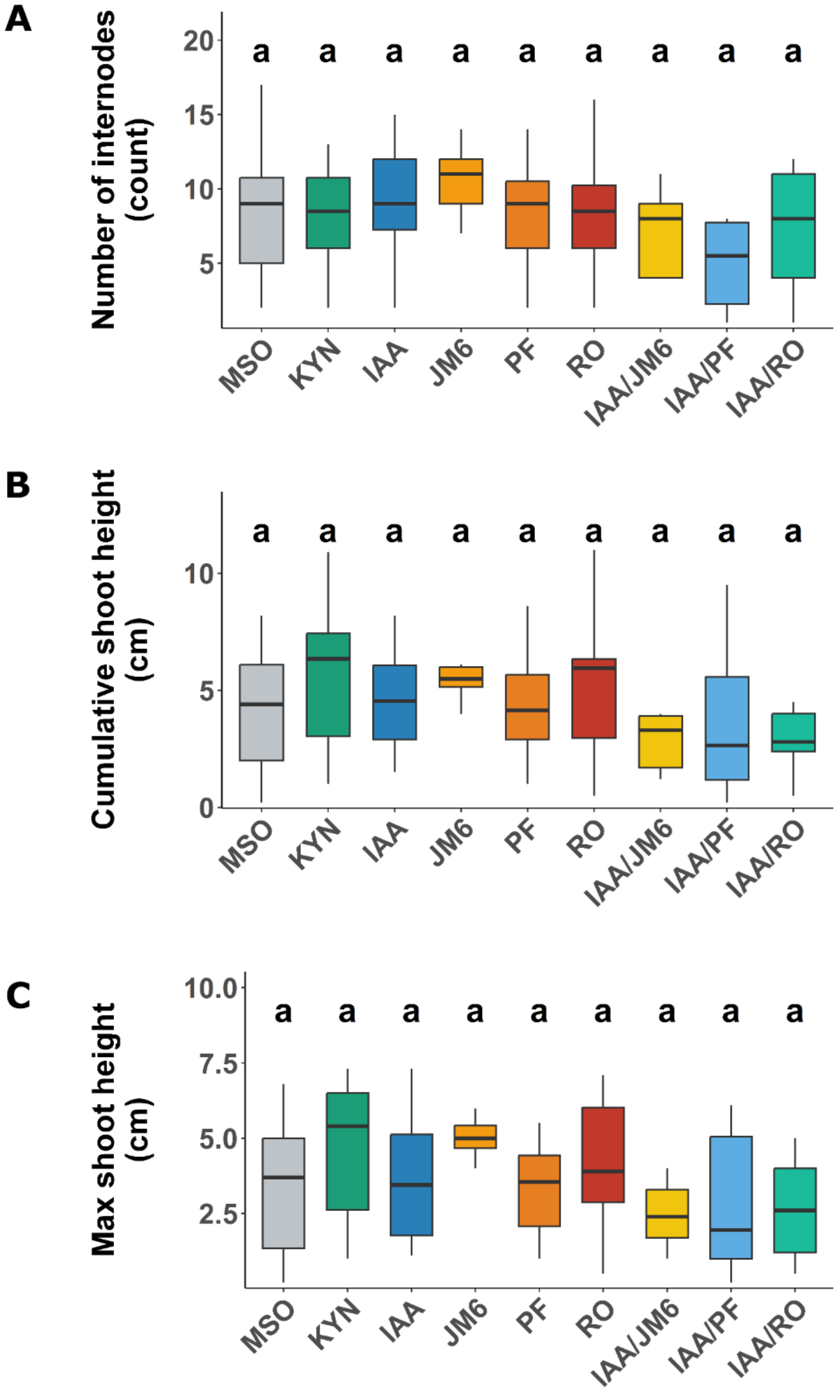

**Figure S3. Effects of KYN and IAA on rooting and shoot growth in *Hypericum perforatum* explants.**

Maximum root length (cm) of rooted explants following treatment with MSO (control), kynurenine (KYN), or indole-3-acetic acid (IAA). (B) Maximum shoot height (cm) comparing rooted and non-rooted explants. (C) Rooting frequency (proportion of explants rooted) for MSO, KYN, and IAA treatments. Bars in (C) represent observed proportions ± 95% Wilson confidence intervals. Statistical comparisons were performed using a binomial generalized linear model for rooting frequency and linear models for continuous traits, with Tukey-adjusted post hoc tests (α = 0.05).

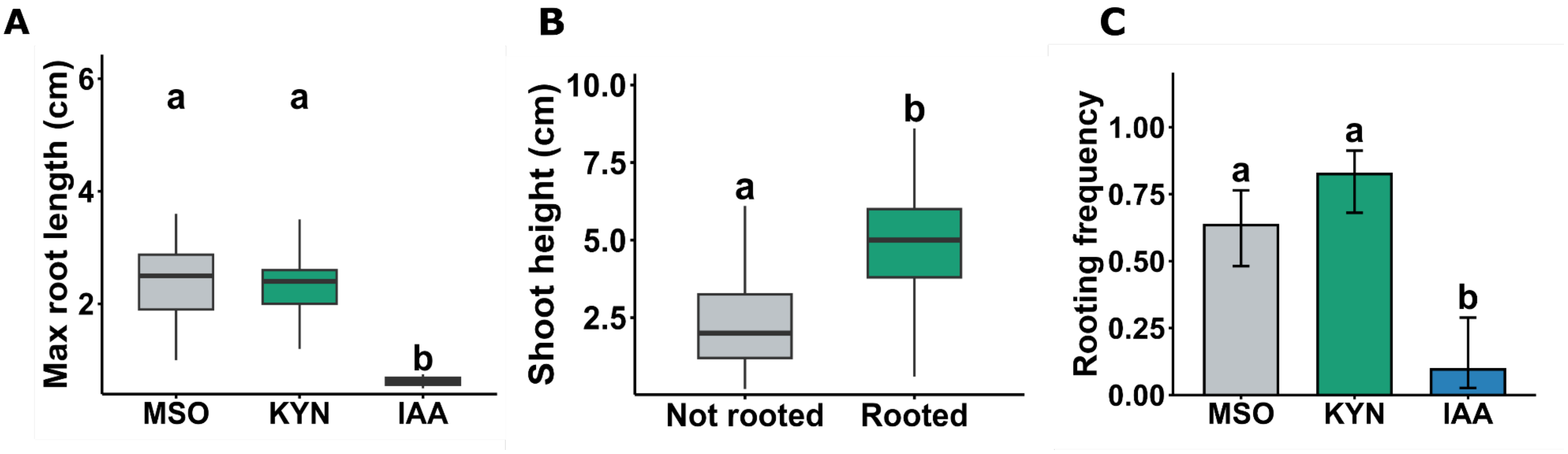
